# ACE Configurator for ELISpot (ACE): Optimizing Combinatorial Design of Pooled ELISpot Assays with an Epitope Similarity Model

**DOI:** 10.1101/2023.09.02.554864

**Authors:** Jin Seok Lee, Dhuvarakesh Karthikeyan, Misha Fini, Benjamin G. Vincent, Alex Rubinsteyn

**Affiliations:** Lineberger Comprehensive Cancer Center, University of North Carolina at Chapel Hill, Chapel Hill, NC; Division of Hematology, Department of Medicine, University of North Carolina at Chapel Hill, Chapel Hill, NC; Department of Microbiology and Immunology, UNC School of Medicine, Chapel Hill, NC, USA; Computational Medicine Program, UNC School of Medicine, Chapel Hill, NC, USA; Curriculum in Bioinformatics and Computational Biology, UNC School of Medicine, Chapel Hill, NC, USA; Department of Genetics, University of North Carolina at Chapel Hill, Chapel Hill, NC 27599 USA

**Keywords:** ELISpot, immunological assay, high-throughput assay design, assay optimization, deconvolution, protein language model

## Abstract

The ELISpot assay is a powerful *in vitro* immunoassay that enables cost-effective quantification of antigen-specific T-cell reactivity. It is widely used in the context of cancer and infectious diseases to validate the immunogenicity of epitopes. While technological advances in hardware and software have kept pace with the need for increased throughput, assay design and deconvolution methodology have largely remained stagnant. Current methods for designing multiplexed ELISpot assays are restricted to preset configurations, lack support for high-throughput scenarios, and ignore peptide identity during pool assignment. We introduce the ACE Configurator for ELISpot (ACE) to address these gaps. ACE generates optimized peptide-pool assignments from highly customizable user inputs and handles positive peptide deconvolution using assay readouts. We present a novel sequence-aware pooling strategy, powered by a fine-tuned ESM-2 deep sequence model to identify immunologically similar peptides, reducing the number of false positives and subsequent confirmatory assays. To validate the performance of ACE using real-world datasets, we conducted a comprehensive benchmark study against various design heuristics, deconvolution methods, and experimental conditions, contextualizing design parameter choices with their impact on precision and number of total pools. Our results demonstrate ACE’s capacity to further increase precision of identified immunogenic peptides, maximizing experimental efficiency at the bench-side. ACE is freely available as an executable with a graphical user interface and command-line interfaces at https://github.com/pirl-unc/ace.

## INTRODUCTION

T-cells play a central role in the immune response against pathogenic infections, malignancies, allergens, or even healthy tissues [1-6]. The process of identifying the specific molecular determinants responsible for a T-cell response, known as *epitope mapping*, is important for vaccine development as well as illuminating the mechanisms underpinning autoimmunity, transplant rejection, anti-tumor activity, and other immunological phenomena [7-10].

The enzyme-linked immunosorbent spot (ELISpot) assay has been widely used for epitope mapping, measuring peptide specific T-cell cytokine secretion, most commonly interferon-gamma (IFN-*γ*) [9, 11-13]. When performed with one peptide per well, epitope mapping through single-plex ELISpot requires increasing efforts, reagents, and biological samples with each additional peptide. In contrast, multiplexed ELISpot enables immunologists to assess the immunogenicity of numerous candidate peptides by combining multiple peptides in the same well [14, 15]. The increased throughput of this approach has been instrumental in identifying immunogenic epitopes in diverse disease contexts such as cancer, HIV, MERS, and SARS CoV-2 [16-27].

Multiplexed ELISpot involves two distinct algorithmic steps: a design method for the allocation of peptides to pools and a deconvolution method for determining immunogenic peptides. Approaches to creating ELISpot designs aim to ensure unique co-occurrence of peptide pairs across replicates to aid deconvolution of pool spot counts. Deconvolution is typically performed using an empirical two-step method: first, by identifying positive pools through activity thresholding, and subsequently conducting an additional round of ELISpot to individually assess all peptides present in those pools [23, 28]. Recently, statistical methods have shown to achieve high accuracy on deconvolution of individual peptide activities [28]. However, adoption of statistical methods has been limited.

Contrary to the widespread adoption of ELISpot, the number of publicly available assay tools is surprisingly limited. DeconvoluteThis [29] utilizes a Monte Carlo method to approximately satisfy the co-occurrence criteria. However, the authors report its performance under a strong prior of expected positivity rate and the software is no longer available. Additionally, Strandberg’s thesis [30] formalizes pooled ELISpot configuration as a Partially Balanced Incomplete Block Design (PBIBDs) [31, 32]. Although Strandberg’s work is accompanied by a webserver tool, it has restrictive assumptions about the assay design such as predetermined numbers of pools and replicates.

Furthermore, the existing tools fundamentally overlook peptide identities when allocating peptides to pools. In practice, many experiments generate libraries where peptides share some nontrivial degree of similarity. Examples include libraries of overlapping peptides from larger antigens and mutational scans that generate related peptides. Recent advances in protein language models such as ProtBERT [33], ProtT5 [33], and ESM [34] have demonstrated a powerful capacity to capture diverse functional features from protein sequence alone. Taken together, we hypothesized that peptide-pool assignments that consider both sequence and functional similarities provide a new dimension to enhance the experimental efficiency of ELISpot, especially in reducing the total number of required pools (sum of first-round and second-round assays) and increasing the precision of identifying immunogenic peptides.

Here, we present ACE (ACE Configurator for ELISpot), a software that facilitates sequence-aware ELISpot assay design and deconvolution of immunogenic peptides. We fine-tuned ESM-2 to predict epitope similarity and demonstrate ACE’s robustness by rigorous benchmarking on various real-world scenarios.

## RESULTS

### Overview of ACE Workflow

ACE supports two main functions that together provide an end-to-end ELISpot experimental optimization: sequence-aware ELISpot assay configuration and deconvolution of positive peptides (Figure 1). These functions are available to the user as a standalone executable with a graphical user interface (GUI) and as a command-line interface (CLI) in a python package.

**Figure 1.**
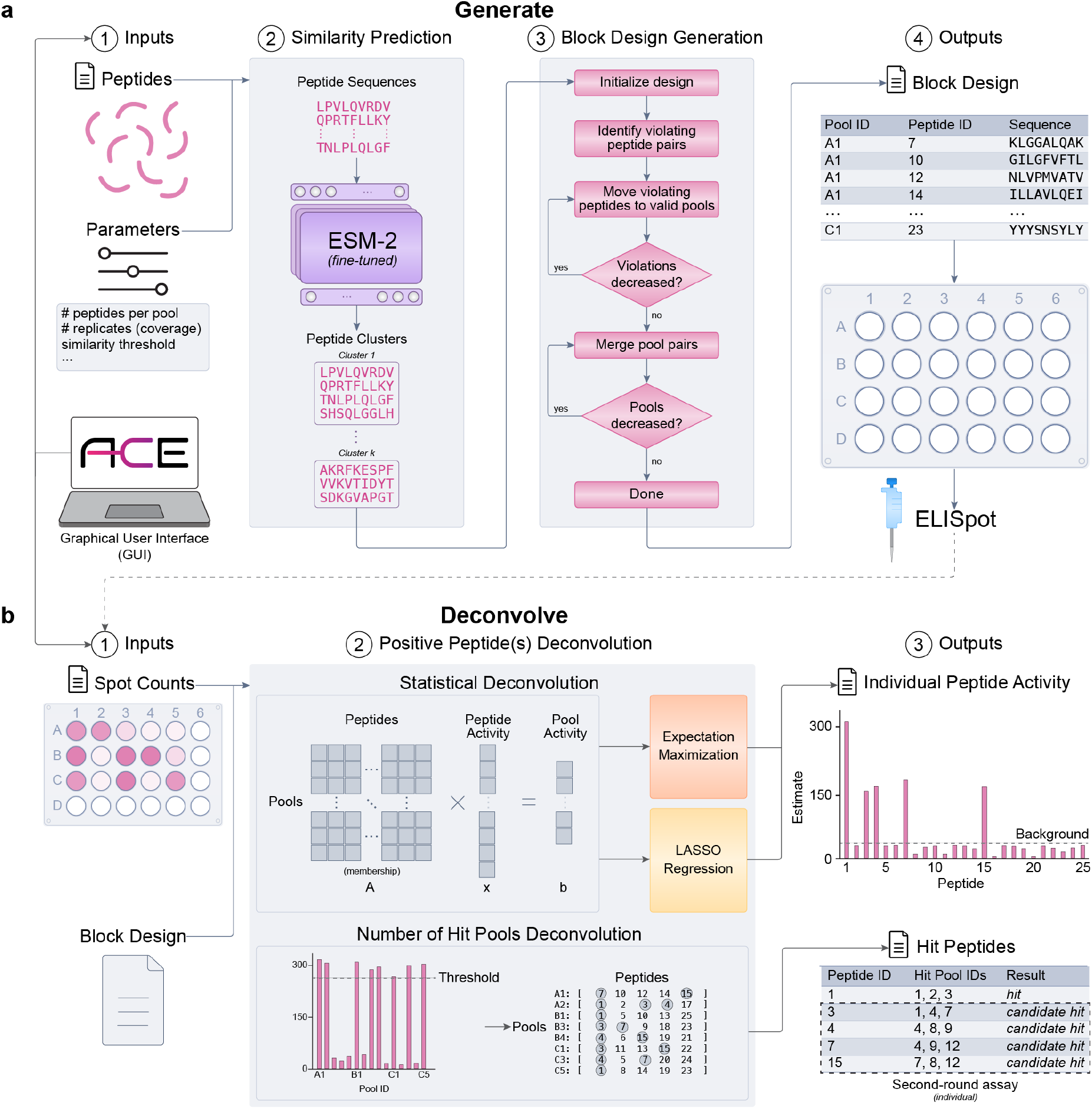
Overview of ACE. **(a)** ACE block design generation algorithm. Inputs include required assay parameters and optional sequence-specific parameters. Pairwise peptide similarity is predicted by the fine-tuned ESM-2 protein language model and grouped into clusters that seed the first coverage of block design. Random swaps are used to reduce the number of peptide co-occurrence violations. The final peptide-pool assignment is mapped onto standardized microtiter plates and output as a tabular spreadsheet for ease of use at the bench. **(b)** ACE immunogenic peptide deconvolution algorithm. The deconvolution module receives ELISpot spot count data and the original ACE block design output. Users can choose either statistical or empirical deconvolution. Statistical deconvolution infers the individual peptide-level spot count while empirical deconvolution relies on a user-supplied positive spot count threshold to binarize immunogenicity.

#### Block Design Generation

ACE’s ‘*generate’* function requires several parameters such as the total number of peptides, the number of desired peptides per pool, and the number of technical replicates (coverage) (Figure 1a). Optional parameters include the peptide sequences, sequence similarity threshold, and the size of microplate (e.g., 96-well plate). To assign peptides to pools, ACE starts from a randomly mapped assignment. ACE then attempts to heuristically transform the random design into a valid PBIBD by iteratively swapping peptides which violate non-overlap constraints. If peptide sequences are supplied, ACE uses its fine-tuned ESM-2 model [35] to compute pairwise similarities between the peptide sequences and groups the pairs that have a similarity greater than the threshold specified. ACE implements this novel sequence similarity approach to ELISpot design because placement of immunogenic peptides significantly affects the experimental efficiency (Supplementary Note 1). Next, using a greedy random swap heuristic that respects the peptide clusters in the first coverage, subsequent coverage peptide-pool assignments are optimized by minimizing the number of violations (peptide pair co-occurrences across replicates). The pooled ELISpot design generation outputs a tabular spreadsheet with peptide IDs, peptide sequences, pool IDs as well as plate and well IDs for usability at the bench-side.

#### Immunogenic Peptide Deconvolution

ACE provides two classes of methods to the user to perform positive peptide identification (Figure 1b). The first is empirical deconvolution of peptides using pool-level positivity criteria to identify peptides from the union of positive pools. In this setting, ACE supports various ‘empirical rules’ for determining immunogenicity [36] and accepts a user-supplied minimum threshold for a pool to be considered positive. Furthermore, ACE distinguishes between two groups of hit peptides: confident hits and candidate hits. Whereas confident hits have at least one positive pool that maps to uniquely to that hit peptide, candidate hits co-occur with other hit peptides across all coverages and require additional assays to ascertain their positivity. We use this distinction in our calculation of an assay’s total number of pools, defined as the sum of the first-round pools plus the number of candidate peptides. The second method offered by ACE is statistical deconvolution that allows the direct inference of peptide-level activities from pooled spot count values. We implement the previously described Expectation Maximization algorithm for deconvolution [28] as well as a Lasso regression method which we selected for its propensity to identify sparse vector solutions [37].

### ACE Block Design Algorithm Outperforms Randomized and Repeated Block Designs

Since existing multiplex ELISpot assay design techniques use some form of PBIBD constraints, we confirmed whether decreasing peptide pair co-occurrences results in more precise detection of immunogenic peptides for a fixed number of pools (Figure 2a). We traced the evolution of repeated block designs, which maximally violate the peptide non-overlap constraints, across iterations of ACE’s design optimizer. We simulated ACE ELISpot configuration generation (without peptide similarity prediction) 100 times with each iteration simulated 100 times (i.e., 10,000 configurations in total) for 400 peptides in groups of 10 with 3x coverage (400/10/3) with 40 positive peptides. In these simulations, we applied random experimental effects to the peptide activity levels. The number of violations was observed at each iteration along with the deconvolution precision and sensitivity. The ACE randomized swap strategy consistently reduced the number of violations over the number of iterations. As expected, we observed a monotonic relationship between the number of violations and deconvolution precision. Notably, the two metrics saturated and exhibited minor oscillations after a sufficiently low number of violations was achieved. We suspect that the fluctuation arises from the shifts in arrangement of immunogenic peptides, which affects the number of false positives (Supplementary Note 1). The sensitivity remained relatively constant, and the minor fluctuations were reflective of the random effects applied in the simulations. These patterns held across various assay sizes and configurations as well, indicating a more general trend reflective of the nature of the problem itself.

**Figure 2.**
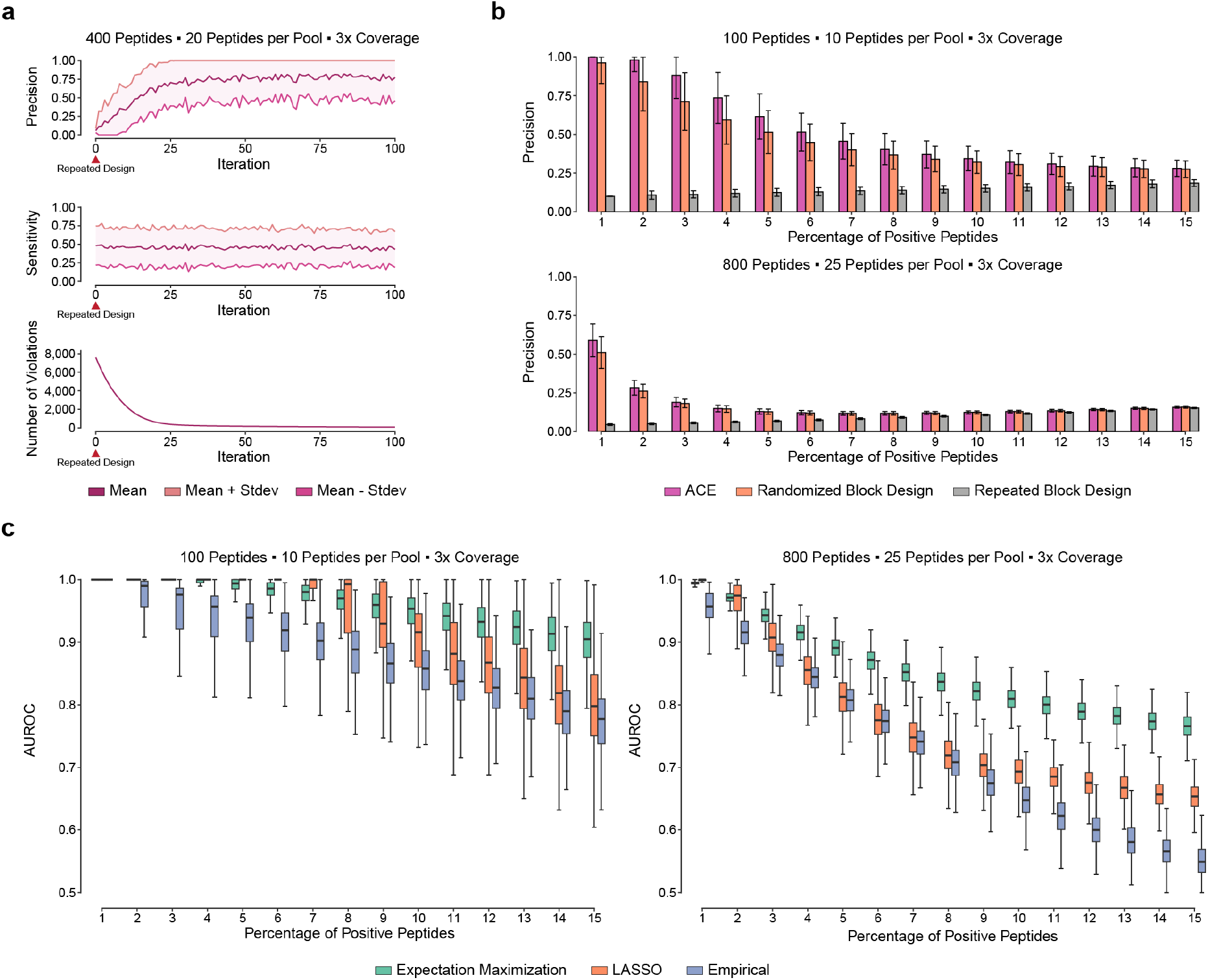
*In silico* evaluation of ACE performance for non-overlapping block design generation and hit peptide deconvolution. **(a)** Relationship among the optimization iterations, precision, recall, and number of violations simulated over 100 optimization trajectories with ±1 standard deviation. **(b)** Precision of single-round deconvolution results for ACE, randomized block design, and repeat block design across various thresholds of peptide positivity in both the 100/10/3 and 800/25/3 experiments. Error bars represent ±1 standard deviation across 1,000 simulations. **(c)** Box and whisker plot showing median and interquartile range of single peptide activity deconvolution AUROC for empirical deconvolution, lasso regression, and expectation maximization.

We next benchmarked the ACE block design generation algorithm against two baseline block design strategies. The first was the repeated block design which we considered as the naïve baseline where peptides were arbitrarily assigned into equally sized groups and the assignment was repeated across all replicates. The second was a randomized block design, where peptides were assigned randomly to pools without replacement in each replicate. While we were unable to identify any studies that employed this strategy, the randomized block design represents a more robust computational baseline.

For the ELISpot configuration with 100 peptides with 10 peptides per pool and 3x coverage (100/10/3), the average first-round deconvolution precision across 1% to 15% positive peptides was 0.529, 0.470, and 0.141 for ACE, randomized design, and repeated design, respectively. The average total pools for this experiment configuration were 53, 54, and 83 for the ACE, randomized design, and repeated design (Figures 2b and S1a). The average precision values across 1% to 15% positive peptides for the 800/25/3 configuration were 0.177, 0.170, and 0.094 for ACE, randomized design, and repeated design, respectively. The average numbers of pools for the configuration were 565, 568, and 722 (Figure S1b). We additionally performed benchmark analysis for ELISpot configurations with 200 peptides and 400 peptides and observed similar trends (Figure S1c-d).

Furthermore, we benchmarked ACE against DeconvoluteThis [29] and found that ACE and DeconvoluteThis performed similarly for all the ELISpot configurations originally reported by Roederer and Koup (Figure S2).

### Comparison of ACE Deconvolution Methods

ACE provides three deconvolution methods for pooled ELISpot data given spot forming units (SFUs) per well: empirical, expectation maximization, and LASSO. We evaluated the area under the Receiver Operating Characteristic curve (AUROC) of first-round deconvolution predictions given simulated ground truths for each method across 100, 200, 400, and 800 peptides (Figure 2c and Figure S3). As the baseline method, we employed the commonly used empirical deconvolution approach to identify putative hit peptides by mapping back the peptide-pool assignment from positive pools [36]. Given the absence of a continuous score for AUROC calculation under empirical deconvolution, we artificially set this score to be the coverage of each hit peptide ranging from 0 to the number of replicates. We evaluated two statistical deconvolution methods against this baseline approach -EM and LASSO regression. The EM algorithm iteratively optimizes for the maximum likelihood vector of individual peptide spot counts given the observed pool spot count, as previously described [28]. For 100 peptides (10 peptides per pool and 3x coverage), the average AUROC values were 0.961, 0.931, and 0.882 for the EM algorithm, LASSO regression, and empirical approach, respectively, across 1% to 15% positive peptide percentages. For 800 peptides (25 peptides per pool and 3x coverage), the average AUROC values were 0.854, 0.768, and 0.724, respectively. The simulation results for 200 and 400 peptides exhibited similar trends in favor of the EM approach (Figure S3). We further note that the general downward trend in the AUROC values with the number of positive peptides is attributable to the increase in the number of positive pools, commensurate with the number of false positives.

### Epitope Similarity Further Optimizes ELISpot Assays in Real-World Scenarios

ACE incorporates a neural engine that leverages a fine-tuned ESM-2 model to facilitate epitope similarity prediction and clustering. The model was trained under a contrastive loss objective, designed to bring peptides that share a common T-cell receptor (TCR) closer in the latent embedding space compared to peptides with no shared TCRs (Supplementary Note 2). Compared to the off-the-shelf ESM-2 model, the fine-tuned model assigns high pairwise similarities to peptides that bind to the same TCR even when they share a large Levenshtein edit distance (Figure S4). Based on the two-sample Kolmogorov Smirnov (KS) tests, the D-statistic values were 0.418, 0.427, and 0.476 for the Levenshtein similarity (*p* = 5.149 ∙ 10^−9^), baseline ESM-2 (*p* = 2.095 ∙ 10^−9^), and finetuned ESM-2 (*p* = 1.088 ∙ 10^−11^). While all the comparisons were statistically significant, we achieved the greatest separation with ESM-2 fine-tuning (Figure 3a-b). To evaluate the effectiveness of the peptide clustering strategy, we tested the feature using four different real-world datasets: held-out IEDB peptides [38], immunogenic SARS CoV-2 peptides [39, 40], and cross-reactive MAGE A3 antigens for adoptive T-cell therapy [41]. Hereafter, we refer to our ACE ELISpot configuration generation approach without peptide similarity prediction as ACE and the approach with the prediction as ACE-S.

**Figure 3.**
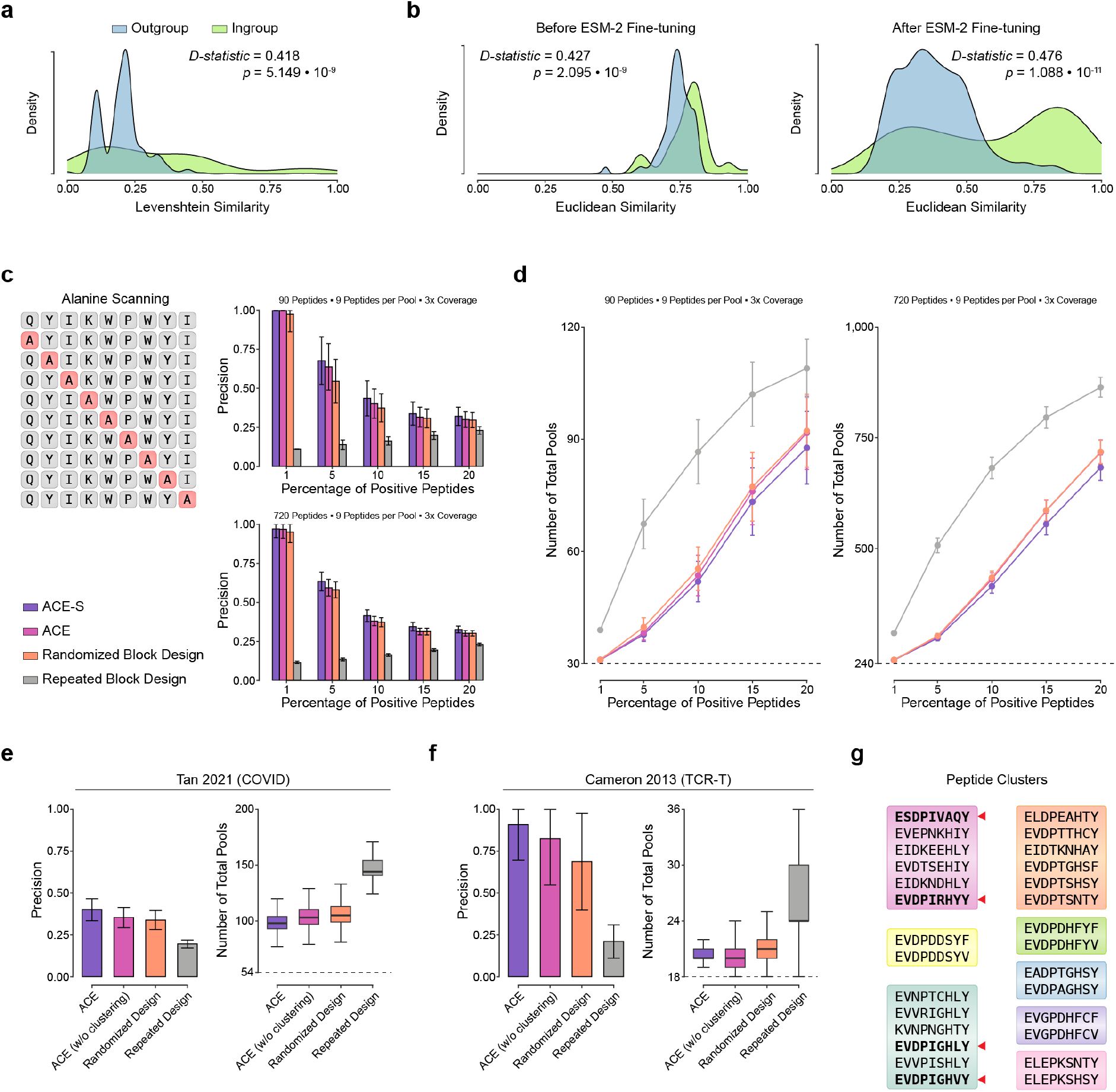
Evaluation of ACE sequence similarity prediction for efficient ELISpot assay in real-world datasets. **(a)** Distribution of pairwise sequence Levenshtein similarity scores stratified by peptides that share TCR specificity (in-group peptides) and those that have no shared common TCRs (out-group peptides). **(b)** Distribution of pairwise Euclidean distances computed on sequence embeddings from in-group and out-group epitopes shown for both off-the-shelf ESM-2 model and its fine-tuned counterpart. **(c)** The left-panel shows a schematic for alanine scan. The right panels show mean precision of simulated alanine-scanned peptide configurations by ACE with and without sequence similarity, randomized block, and repeat block design across various positivity levels. Error bars indicate ±1 standard deviation. **(d)** Line plots of total pools (first-round pools plus candidate hit validation pools). **(e)** Evaluation of ACE sequence similarity module on held-out COVID data (*n* = 177, *n*_*immunogenic*_ = 37). **(f)** Evaluation of ACE in TCR cross-reactivity setting, clustering peptides predicted to bind MAGE A3 TCR. **(g)** Clusters from MAGE A3 experiment with immunogenic peptides bolded and highlighted with red arrows.

Mutational scans substitute each residue of a peptide sequence and test for loss of the property of interest, such as immunogenicity. The alanine scan is a type of mutational scan where each amino acid along a protein is swapped specifically for alanine to assess critical residue positions (Figure 3c) [42]. In studies of T and B-cell reactivity, alanine scans and other mutational scans have been used to investigate which epitope positions are significant for cognate receptor recognition [43, 44]. We benchmarked ACE’s sequence similarity function in the context of testing immunogenicity for alanine scanned 9-mers from the IEDB database [38]. We randomly selected 10, 20, 40, and 80 starting nonamers, which resulted in 90, 180, 360, and 720 alanine-scanned peptides. In the *n* = 90 scenario (9 peptides per pool with 3x coverage), the average precision values across the positive peptide percentages were 0.554, 0.531, 0.498, and 0.168 for ACE-S, ACE, randomized block design, and repeated block design, respectively. In the *n* = 720 scenario (9 peptides per pool with 3x coverage), the average precision values were 0.538, 0.512, 504, and 0.168 for ACE-S, ACE, randomized block design, and repeated block design, respectively. The sequence-aware ACE-S also outperformed the other three approaches for *n* = 180 and *n* = 360 configurations (Figure S5). Consistent with the benchmark experiment studies so far, the precision inversely correlated with the number of total pools (Figure 3d).

While the added benefit in the alanine-scanning evaluation was marginal, we maintain that the above stands as an evaluation of both the strategy of clustering similar peptides and the ability to accurately perform this clustering. To test the effect of the peptide clustering strategy alone, we performed the same experiment on alanine-scanned peptide sequences with starting sequences seen during training. Sequence-aware clustering by ACE-S on these explicitly trained starting sequences resulted in precision values of 0.602 and 0.622 for *n* = 90 and *n* = 720 scenarios, respectively (Figure S6) demonstrating the efficacy of peptide similarity pooling, independent of peptide similarity calculation.

We next evaluated the effect of applying sequence-based clustering on ELISpot experimental efficiency on real-world datasets where immunogenicity was validated *in vitro*. The first was a set of curated SARS-CoV-2 epitopes assessed for cross-reactivity to high precursor TCRs [39]. This dataset consisted of 177 known CD8+ and CD4+ epitopes. We considered the 22 CD8+ epitopes to be immunogenic and the 155 CD4+ epitopes to be non-immunogenic, and simulated ELISpot designs pooling 10 peptides per pool and 3 technical replicates. For this dataset, the average first-round deconvolution precision values were 0.400, 0.354, 0.344, and 0.198 for ACE-S, ACE, randomized block design, and repeated block design, respectively (Figure 3e).

We further examined if sequence similarity pooling would be helpful in large protein-level epitope scans. We identified 1,265 nonamers over the entire SARS-CoV-2 spike protein sequence by sliding window (k = 9) and found 105 (8%) matching immunogenic nonamers from the ImmuneCODE database with known TCR binding [40] (Figure S7). We found that ACE-S and ACE performed similarly in this sequence set, due to the inherent sequence similarity arising from applying a sliding window. Although there was no significant benefit to clustering, we note that even in adversarial settings, ACE-S performs on par with ACE and better than the randomized design heuristic.

Finally, we experimented with a set of 37 epitopes tested for off-target T-cell response due to cross-reactivity in a MAGE A3-directed adoptive T cell therapy [41]. We used this dataset to evaluate the ability of our peptide similarity prediction module to group peptides that bind to the same TCR. While not perfectly grouped, we found that the partial grouping of the MAGE A3 reactive epitopes resulted in first-round deconvolution precision values of 0.909, 0.824, 0.686, and 0.211 for ACE-S, ACE, randomized block design, and repeated block design, respectively (Figure 3f-g).

We conclude the analysis of the ACE’s sequence-aware approach with the following recommendation. Its integration has the potential to significantly enhance the positive peptides precision while performing just as well as no sequence clustering at the worst case. Given the marginal computational cost incurred by the model during inference, and the added benefit of its inclusion we set the ACE default to factor epitope similarity.

### Exploration of ELISpot configuration design space

We further examined ACE and the multiplexed ELISpot analysis design problem with an exploration of the parameter space and its impact on performance. The optimal configuration for a fixed peptide library depends largely on the experimentalists’ constraints. Common considerations to optimize for are experimental readout time, reagent usage, and manpower. Time in an ELISpot assay depends on whether second-round assays are required. As a proxy, we measure first-round precision. The second consideration is the total pool count, which serves as a direct proxy for the amount of reagents consumed over the entire experiment. Interestingly, varying the peptides per pool and coverage across different number of total peptides resulted in relatively constant heatmaps for both the precision and number of total pools, relative to the number of starting pools. We identified two different optimal regions for the ELISpot design: maximal precision occurs with a low number of peptides per pool and a high number of replicates whereas minimal total number of pools occurs with an intermediate number of peptides per pool and low coverage (Figure 4).

**Figure 4.**
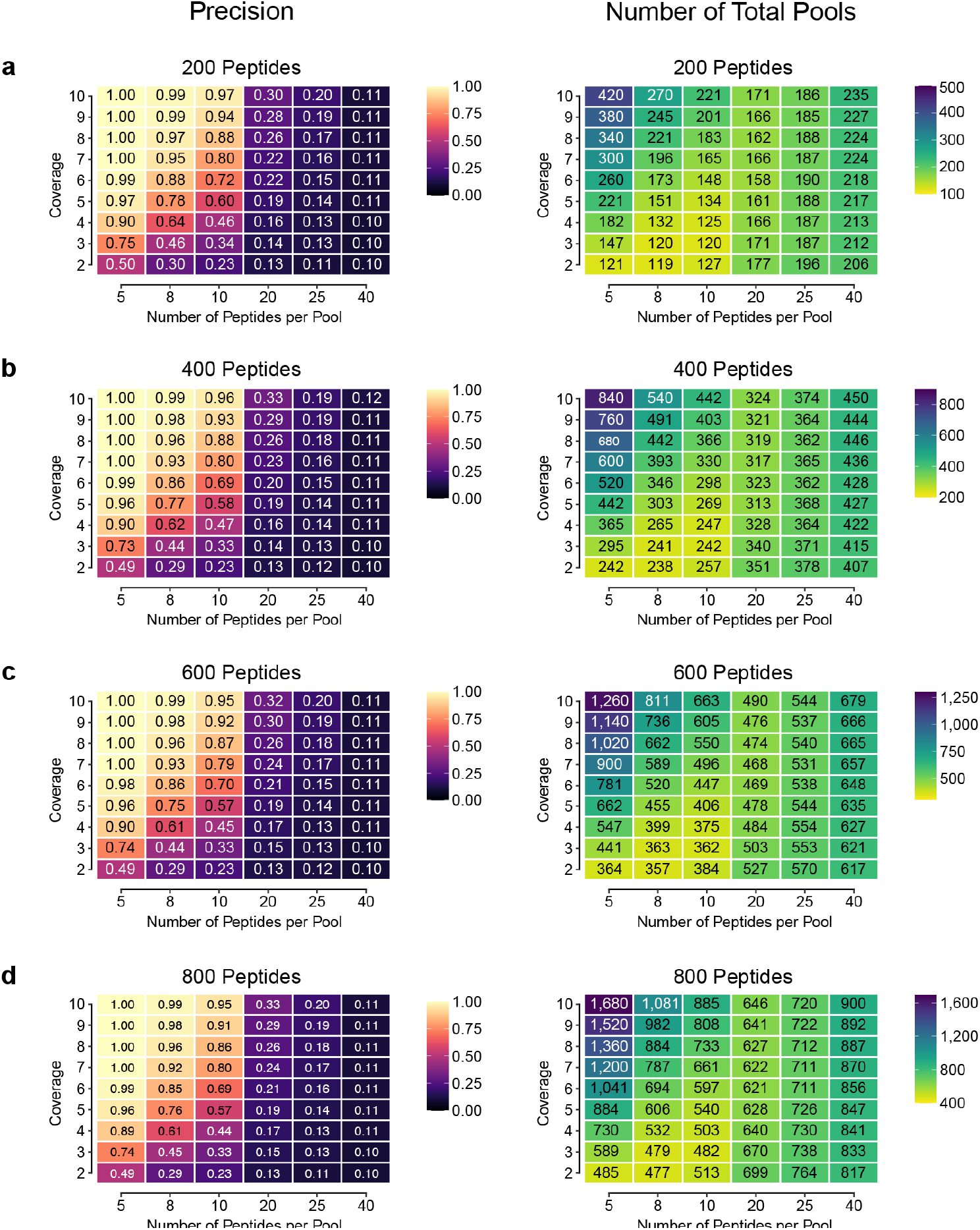
Exploration of ELISpot design optimization space. Parameter sweeps across the coverage and pool size for a fixed number of peptides were performed with precision and total number of pools measured. **(a-d)** 200, 400, 600, and 800 peptides were simulated for 1,000 runs under a 10% positivity rate and averages are shown within the boxes. Color bars are scaled locally using the relative minimum and maximum for each heatmap.

### ACE algorithm runtime and peak memory usage scale linearly with the user inputs

We assessed ACE’s scalability by analyzing runtime and peak memory usage. We explored three scalability axes: total peptide count, peptides per pool, and design coverage. Through simulations, we measured runtime and peak memory usage for ACE’s *‘generate’* while varying these parameters. Fixing peptides per pool at 10 and coverage at three, runtime and memory showed a linear correlation with peptide count (Figure 5a). With a fixed peptide count of 1,000 and varied peptides per pool (10 to 30), runtime exhibited modest fluctuations (Figure 5b), reflecting ACE’s preset iterations. Altering technical replicates (3 to 10) for 1,000 peptides in 10-pool settings showed linear runtime trends, but peak memory usage did not follow this pattern (Figure 5c). At *coverage* = 10, peak memory surpassed usage at *coverage* = 3 by over 10x, suggesting polynomial or exponential scaling. ACE’s memory usage remains manageable on typical laptops.

**Figure 5.**
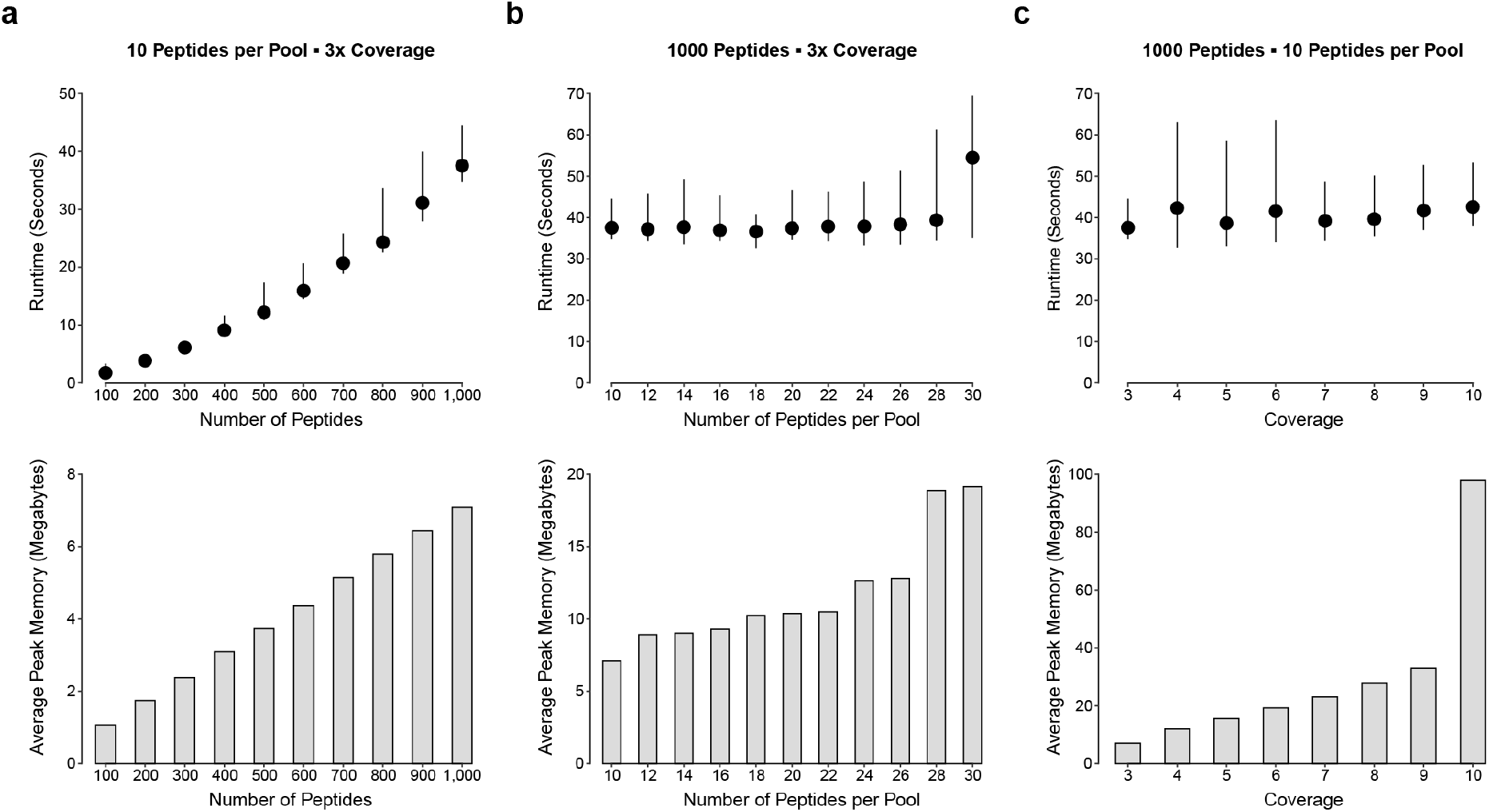
Scaling ACE. ACE *‘generate’* function peak memory usage and runtime measured across various parameter ranges with epitope similarity inference. **(a)** Runtime and peak memory usage with varying numbers of peptides while fixing 10 peptides per pool and 3 technical replicates. **(b)** Runtime and peak memory usage across different numbers of peptides per pool for block designs with 1,000 peptides and 3 technical replicates. **(c)** Runtime and peak memory usage across different numbers of technical replicates for block designs with 1,000 peptides and 10 peptides per pool.

## DISCUSSION

Detection of antigen-specific T-cell responses is fundamental to our ability to study the adaptive immune system, providing the context to unravel higher-order phenomena such as cross-reactivity and immunodominance. While multiplexed ELISpot offers an experimental platform to perform epitope mapping on thousands of peptides, existing design generation and deconvolution methods are limited in flexibility or usability and do not fully exploit the inherent biological sequence properties in their design. In response to these limitations, we developed the ACE Configurator for ELISpot (ACE), which leverages advances in deep learning methods in sequence modeling to offer a novel sequence-aware arbitration strategy that enhances assay efficiency and precision.

To demonstrate ACE’s robustness, we benchmarked its block design generation and immunogenic peptide deconvolution approaches using a carefully constructed simulation framework, with further validation using real-world datasets. Our analysis demonstrated the effectiveness of the ACE core algorithm, particularly highlighting the added benefit of sequence similarity in various experimental settings.

In practical application, we expect ACE’s sequence similarity prediction feature will be especially useful in experimental designs that investigate related peptide epitopes. These contexts include evaluation of cross-reactivity across a set of antigens for the same TCR clonotype. We demonstrated this by using the *in vitro* validated cross-reactive MAGE A3 antigens in the context of adoptive T-cell therapy. We have shown that the cross-reactive immunogenic peptides are successfully clustered and pooled together by ACE. Although this antigen set was small (*n* = 37), this trend held with larger peptide libraries across a diverse set of sequences. Moreover, our analysis using trained sequences in the alanine scan demonstrates that the model produces improved embeddings for sequences closer to available training data. As such, the ACE neural engine is poised to improve over time as more diverse data are integrated into training. We acknowledge the use of peptide similarity as a surrogate for immunogenicity prediction, and while it showcased a meaningful impact, we foresee greater efficacy by combining the peptide similarity module with weak immunogenicity prediction in future iterations of ACE.

Building upon previous statistical deconvolution approaches, we consistently achieve higher AUROC values compared to the standard empirical deconvolution method. However, practitioners seldom use this powerful method. We hope that our transparent evaluation of these methods, coupled with the accessible implementation of empirical and statistical deconvolution in the ACE GUI, will encourage broader adoption.

ACE’s sequence-aware approach offers a promising avenue for advancing ELISpot experimental design. By bridging the gap between sequence-level properties and assay configuration, ACE equips researchers with a powerful tool to unlock the full potential of multiplexed ELISpot in unraveling the complexities of adaptive immune responses. As the field of precision immune medicine continues to expand, the exploration of the ELISpot problem space and efforts in optimization become more crucial. We release ACE and its accompanying analyses and recommendations to better inform pooled ELISpot experimental design, in anticipation of its increased usage in the precision immune medicine space.

## METHODS

### Block Design Generation

ACE is designed to assist in generation of custom configurations and deconvolution of immunogenic peptides. Given the required parameters such as total number of peptides (N), number of peptides per pool (p), and number of replicates (r), ACE initializes a random peptide-pool mapping. It then iteratively minimizes the number of violations by randomly swapping the violating peptides, pushing violations to further replicates. This method shares resemblance to DeconvoluteThis [29] and the random search method [30]. A distinguishing feature between ACE and the existing methods is the clustering of peptides based on predicted TCR reactivity. Clusters of similar peptides inform the first replicate assignment. ACE attempts to minimize violations in subsequent replicates. While faster methods for generating NODs exist that leverage orthogonality in matrix designs, we implement the random-swap strategy as it better facilitates the inclusion of the epitope similarity module.

### Peptide Similarity Clustering

To boost precision and sensitivity by grouping similar peptides that bind to the same TCR, we chose deep sequence models which have outperformed conventional metrics such as Levenshtein and Hamming distances. Coupled with our need for a strong model that could fit on a laptop memory, we opted for the smallest ESM-2 model with six layers and roughly 8 million parameters, downloaded from the HuggingFace repository. We fine-tuned the model using the triplet loss to push embeddings from peptides that share a TCR closer while pulling away embeddings from epitopes that share no TCR (Supplementary Note 2). The pairwise Euclidean similarity scores are computed across all pairs in an assay. Pairs above the sequence similarity threshold are filtered and the top-2 most similar peptides are retained per peptide. Peptide pairs are subsequently clustered by the transitive property. We found this method to outperform other clustering methods such as k-nearest neighbor and agglomerative clustering.

### Simulation

All our benchmark studies were performed using a unified three-step simulation framework. First, we took a specified configuration consisting of assay parameters such as number of peptides, number of true positives, number of peptides per pool, and coverage. Second, we sampled from held-out sequences and immunogenicity labels taken from the IEDB database [38]. Third, we simulated the pool spot count using a bottom-up approach, adding together peptide spot counts per replicate. Peptide spot counts were simulated without stochastic noise to measure design performance and with random effects to evaluate deconvolution methods.

We sampled the peptide spot count from a generalized negative-binomial distribution, as previously described [28, 30] for its application in count data where the variance is greater than the mean [45]. We include as a hyperparameter, the dispersion factor with a default set to (*ϕ* = 1), which is a Poisson distribution. Per-peptide spot count was simulated using a conditional negative binomial conditioned on the immunogenicity status of the peptide. We kept the variance the same for immunogenic and non-immunogenic peptides to best model the variability of precursor TCR frequencies.

To evaluate our simulations, we measured AUROC as a threshold independent method of interrogating the configuration’s unbiased ability to separate the positive and negative peptides, independent of the positivity selection criterion applied (i.e., empirical vs statistical). We also captured practical evaluation metrics such as precision, recall, and F1 score. We considered empirical peptide positivity when peptides occurred in at least *k* positive pools. For all simulations, *k* was set to the number of technical replicates.

### Alanine Scanning Experiment

Alanine scan mutagenesis was selected for its simplicity and reduced peptide state space. Predetermined ELISpot parameters were chosen such as the number of starting peptides, peptides per pool, total replicates, and positive peptide count. Sequences were then sampled from the held-out validation set of unseen epitopes in each run and underwent *in silico* mutagenesis to generate a library of wild-type and mutant peptide sequences. Although our simulation assumed no change in immunogenicity status after undergoing alanine scanning (a necessary simplification given the complexity of TCR binding [45,48]), we believe this assumption effectively conveys the benefit of clustering based on epitope similarity without misrepresenting performance.

### Deconvolution

To identify the peptides that we classify as true positives, several positivity selection criteria exist. Earlier works describe the difficulty in establishing universal cutoffs for positivity and instead discuss “empirical rules” and “statistical tests” that make use of negative wells with reagent-peptides to compare experimental wells [35]. There are three commonly used approaches in the empirical rules setting: minimum threshold for spot counts normalized by cell count, minimum threshold for spot counts as a fold difference compared to the negative controls, and normalization of spot counts via background correction. In the statistical setting, T-tests, Wilcoxon rank sum tests, and permutation tests have been used to compute significance levels between the spot counts of the mock wells and each of the experimental wells to determine positive wells. These methods all rely on reverse mapping of positive pools to identify the potentially positive peptides that are at the union of all the positive pools and occur at a given number of replicates. Alternatively, previous work performs single-shot deconvolution of peptide spot counts from pooled spot counts to separate immunogenic peptides from non-immunogenic ones by approximating solutions to the linear system [28]:

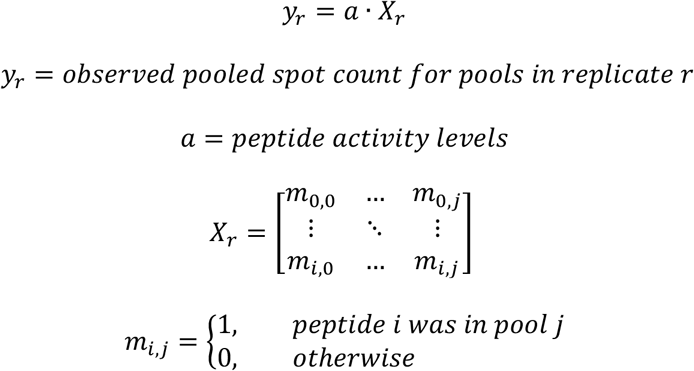

We implemented both statistical and empirical positivity selection criteria and compared these methods against each other. We differentiate between ‘candidate hit’ and ‘confident hit’ to refer to two classes of peptides that are both deemed positive. The candidate hits require additional round of assay to validate immunogenicity.

## ACKNOWLEDGEMENT

We thank Geddes Levenson for feedback and technical help on our figures. We thank Julia Webb for feedback and help with proofreading our manuscript.

## CODE AVAILABILITY

Both the standalone executable with a graphical user interface and the command-line interface with a python package of ACE are freely available at https://github.com/pirl-unc/ace. The benchmark study scripts are available at https://github.com/pirl-unc/ace-analysis.

